# A Comprehensive Enumeration of the Human Proteostasis Network. 1. Components of Translation, Protein Folding, and Organelle-Specific Systems

**DOI:** 10.1101/2022.08.30.505920

**Authors:** The Proteostasis Consortium, Overall coordination, Suzanne Elsasser, Lisa P. Elia, Richard I. Morimoto, Evan T. Powers, Harvard Medical School group (analysis), Daniel Finley, University of California, San Francisco and Gladstone Institutes group I (chaperones, analysis), Eric Mockler, Leandro Lima, Steve Finkbeiner, University of California, San Francisco group II (chaperones, analysis), Jason E. Gestwicki, Northwestern University group (chaperones, analysis), Thomas Stoeger, Kedi Cao, The Scripps Research Institute group (chaperones, endoplasmic reticulum proteostasis, mitochondrial proteostasis, analysis), Dan Garza, Jeffery W. Kelly, Stanford University group (chaperones, translation, mitochondrial proteostasis), Miranda Collier, T. Kelly Rainbolt, Shuhei Taguwa, Ching-Chieh Chou, Ranen Aviner, Natália Barbosa, Fabián Morales-Polanco, Vincent B. Masto, Judith Frydman

## Abstract

The condition of having a healthy, functional proteome is known as protein homeostasis, or proteostasis. Establishing and maintaining proteostasis is the province of the proteostasis network, approximately 2,500 genes that regulate protein synthesis, folding, localization, and degradation. The proteostasis network is a fundamental entity in biology with direct relevance to many diseases of protein conformation. However, it is not well defined or annotated, which hinders its functional characterization in health and disease. In this series of manuscripts, we aim to operationally define the human proteostasis network by providing a comprehensive, annotated list of its components. Here, we provide a curated list of 959 unique genes that comprise the protein synthesis machinery, chaperones, folding enzymes, systems for trafficking proteins into and out of organelles, and organelle-specific degradation systems. In subsequent manuscripts, we will delineate the human autophagy-lysosome pathway, the ubiquitin-proteasome system, and the proteostasis networks of model organisms.

## Introduction

The process by which nascent proteins fold into their native structures co- or post-translationally is imperfect and can fail (Balch et al., 2008; Chiti and Dobson, 2017; Clark, 2004; Gershenson et al., 2014; Powers and Gierasch, 2021; Sala et al., 2017). These failures leave proteins in non-native states that are nonfunctional and potentially toxic, especially if they form aggregates that alter the stoichiometry of macromolecular complexes or interfere with fundamental cellular processes (Balch et al., 2008; Chiti and Dobson, 2017; Chung et al., 2018; Hartl et al., 2011; Knowles et al., 2014). To maintain the proteome in a healthy, functional state—a condition known as protein homeostasis, or proteostasis—all organisms, beginning with the last universal common ancestor (Balch et al., 2008; Draceni and Pechmann, 2019; Powers and Balch, 2013), have cellular components that manage the synthesis, folding, trafficking, and degradation of proteins. These components are collectively designated the proteostasis network (Balch et al., 2008; Jayaraj et al., 2020; Powers and Gierasch, 2021; Powers et al., 2009; Sala et al., 2017). The proteostasis network, while widely discussed in the literature, remains loosely defined because there is no systematic and comprehensive list of its components. The efforts made toward this end have mostly focused on chaperones (Brehme et al., 2014; Shemesh et al., 2021), with less effort devoted to other equally important arms of the proteostasis network involved in transport and degradation. The widespread interest in diseases characterized by failures of proteostasis— especially aging-related neurodegenerative diseases (Hipp et al., 2019; Kaushik and Cuervo, 2015; Labbadia and Morimoto, 2015; Lopez-Otin et al., 2013; Sonninen et al., 2020)—and intensive efforts to identify critical limiting targets for therapeutic interventions suggest that having such a comprehensive description of this network would be invaluable. We address this gap in our knowledge in this and subsequent papers in this series by presenting an enumeration of the human proteostasis network. In this paper, we focus on chaperones, folding enzymes, systems for trafficking proteins into and out of organelles, and the machinery of protein synthesis. Subsequent work will present the annotation of two large and complex systems for protein degradation in eukaryotes, namely the autophagy-lysosome pathway (ALP) and the ubiquitin-proteasome system (UPS), followed by the proteostasis networks of model organisms.

## Results

### Categorization of proteostasis network components

The proteostasis network consists of the systems that regulate and manage protein synthesis, folding, localization, and degradation (Balch et al., 2008; Jayaraj et al., 2020; Powers and Gierasch, 2021; Powers et al., 2009; Sala et al., 2017). The components of protein synthesis include the ribosome and ribosome biogenesis factors; translation initiation, elongation, and termination factors; and the ribosomal quality control (RQC) machinery. The components of protein folding include the canonical chaperone systems (the HSP70s, HSP90s, small HSPs, and TRiC/CCT, and their associated co-chaperones); the folding enzymes (peptidyl-prolyl isomerases and protein disulfide isomerases), some of which also function as co-chaperones; the glycoprotein folding machinery of the endoplasmic reticulum (ER), ranging from oligosaccharyl transferase to the lectin chaperones to the folding-state-responsive glycan trimming enzymes; and other, more substrate- or organelle-specific folding systems (e.g., for collagen). The components of protein localization include the channels through which proteins are shuttled into and out of organelles and the components that recognize substrates for transport. Finally, the components of degradation include the organelle-specific degradation systems in the ER (ER associated degradation, or ERAD) and mitochondria (mitochondrial proteases), the ALP and the UPS, but as noted above the ALP and the UPS will be covered in subsequent papers in the series. We have excluded processes that are essential for protein biogenesis and transport but with indirect influences on proteostasis like, for example, amino acid synthesis and core metabolism (which produces amino acids for protein synthesis), gene transcription and mRNA quality control (which influence proteome composition), and cytoskeletal transport (which contributes to protein localization).

For the proteostasis network, we established “entity-based” and “domain-based” criteria to determine whether a given component should be included in the proteostasis network. The former refers to inclusion of an “entity” (a protein or non-coding RNA) based on positive biochemical, cell biological, or genetic evidence in the literature that it functions in proteostasis. The latter refers to inclusion based on a protein containing a structural domain that is strongly associated with proteostasis; for example, the J-domain, which characterizes cochaperones in the HSP70 system (Kampinga and Craig, 2010). Preliminary lists of proteostasis network components were generated based on reviews and other literature on the relevant systems (see Methods). These preliminary lists were then vetted gene by gene by the authors. For borderline cases, rationales for inclusion or exclusion were presented by Proteostasis Consortium members with subject area expertise and a decision was made based on the consensus of the group.

To organize the proteostasis network components, we sought an annotation system that would convey at a glance a component’s localization and function in proteostasis. Such an approach complements other formalized annotation systems that rely on structured vocabularies, such as the Gene Ontology (Ashburner et al., 2000; Consortium, 2021). Thus, we developed a simple taxonomic scheme consisting of five levels: Branch, Class, Group, Type, and Subtype. We found that five levels were sufficient to convey a general sense of each component’s localization and function while minimizing the number of descriptors. The broadest category, Branch, refers to a component’s localization or membership in an overarching pathway. We defined eight Branch categories: cytonuclear proteostasis, ER proteostasis, mitochondrial proteostasis, nuclear proteostasis, cytosolic translation, proteostasis regulation, the ALP, and the UPS (Figure 1). Note that the term “cytonuclear” refers to components that support proteostasis in both the cytosol and the nucleus, whereas “nuclear” refers to components that primarily support nuclear proteostasis (e.g., histone chaperones). Also note that “proteostasis regulation” refers to components that either control transcription of proteostasis network components genes or control translation as part of, for example, a cell stress response. Class refers to a component’s function in proteostasis (e.g., chaperones, protein transport, etc.), while Group, Type, and Subtype provide increasingly specific descriptors of proteostasis functions within a Class. Our goal was to use only as many descriptors as are minimally necessary to give a basic understanding of a component’s role in proteostasis. Thus, not every component has Type or Subtype annotations. Also, some components have multiple roles in the proteostasis network. These are given multiple entries in our list to reflect each separate role.

**Figure 1.**
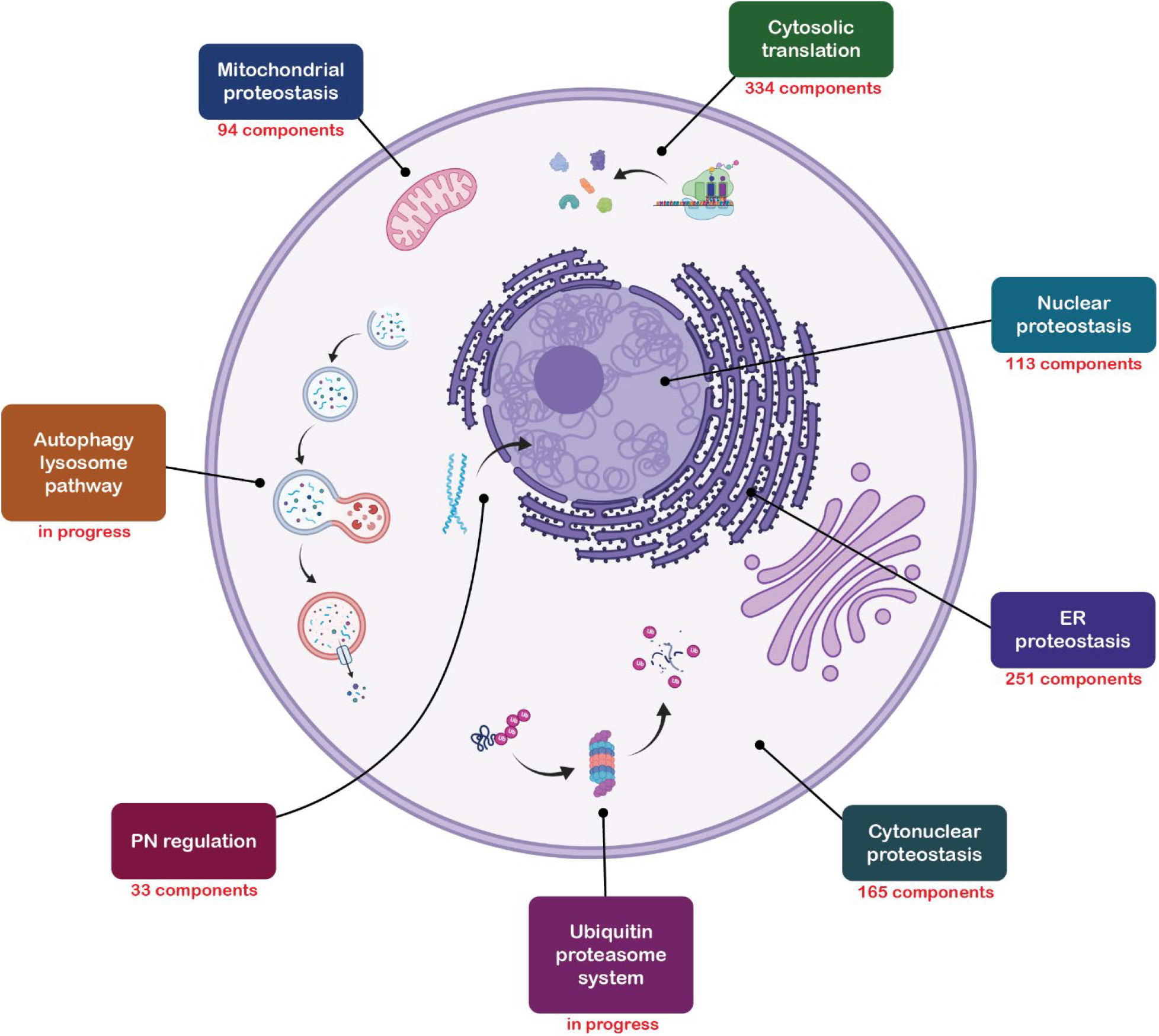
The proteostasis network. The branches of the proteostasis network in our taxonomic system are shown in boxes. The number of components is shown for the six branches covered in this manuscript.

As an illustrative example, we applied our taxonomic scheme to BAG1 (BAG cochaperone 1), a cytonuclear BAG-domain-containing nucleotide exchange factor for HSP70. As shown in Figure 2, this yielded the following annotations: Branch = cytonuclear proteostasis; Class = chaperones; Group = HSP70 system; Type = HSP70 nucleotide exchange factor; Subtype = BAG domain family. Note that annotations are in general not unique to individual proteostasis network components. For example, the five remaining BAG-domain-containing nucleotide exchange factors in humans (BAG2-BAG6) all have the same set of annotations as BAG1 (Figure 2). It is also noteworthy that BAG6 has multiple entries because it is both an HSP70 nucleotide exchange factor and part of a complex involved in the GET pathway for insertion of tail anchored proteins into the ER membrane (Borgese et al., 2019).

**Figure 2.**
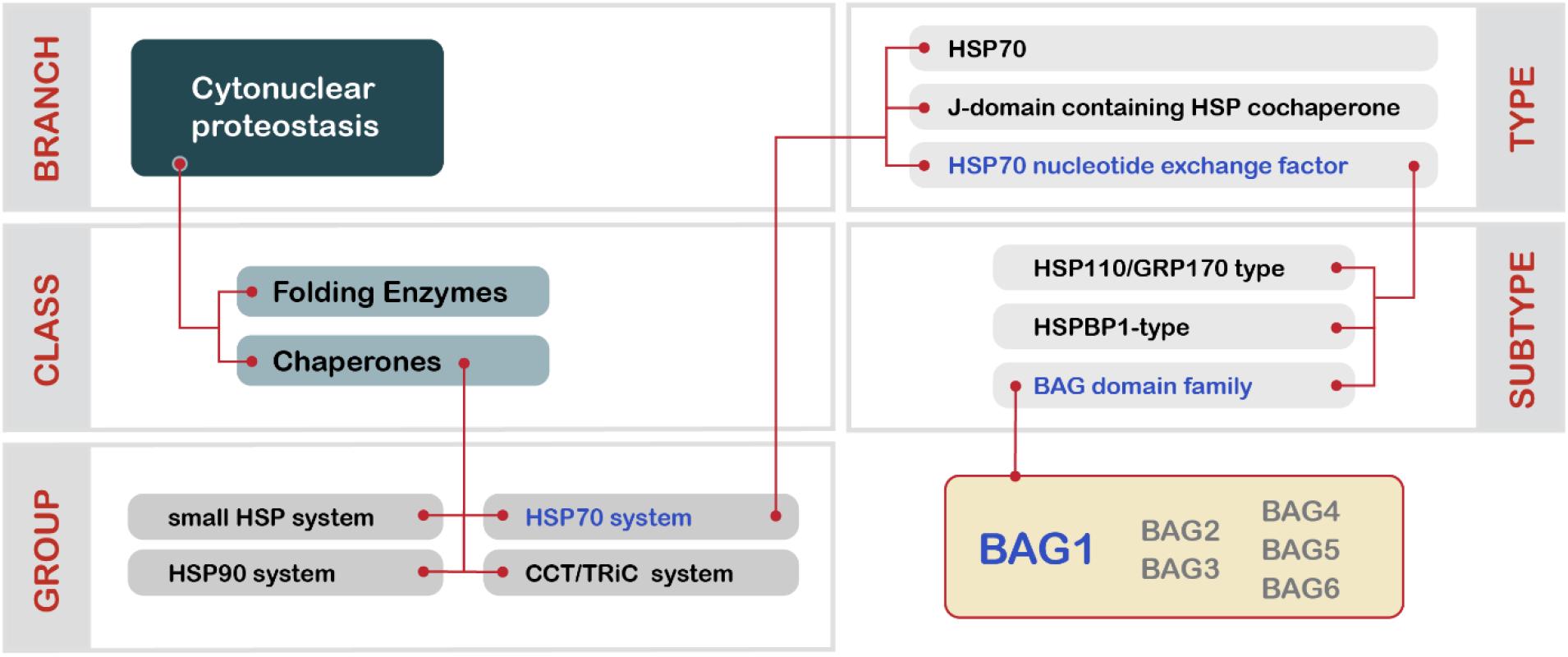
Illustration of the five-level taxonomic scheme applied to proteostasis network components herein using the HSP70 nucleotide exchange factor BAG1 as an example.

### The proteostasis network, excluding the degradative ALP and UPS

The list of proteostasis network components, less the ALP and the UPS, is presented in Supplemental Table 1. There are 1,014 entries in this subset of the proteostasis network representing 959 unique components. Of these, 906 have one entry, 52 have two entries, and one (valsolin containing protein) has four entries. These entries are divided among the Branches shown in Figure 1. All 1,014 entries have Class annotations, divided among 16 categories. Four of these Class categories are used in multiple Branches. For example, the Class annotation “chaperone” applies in the cytonuclear, nuclear, ER, and mitochondrial Branches. This multiplicity reflects the presence of distinct complements of chaperones that have analogous functions in the subcellular compartments in which they reside. The Class annotations “folding enzyme”, “protein transport”, and “organelle-specific protein degradation” also appear in multiple branches and likewise represent sets of components with corresponding proteostasis functions in different compartments. The distribution of Classes among the Branches and the number of components in each Class is illustrated in Figure 3.

**Figure 3.**
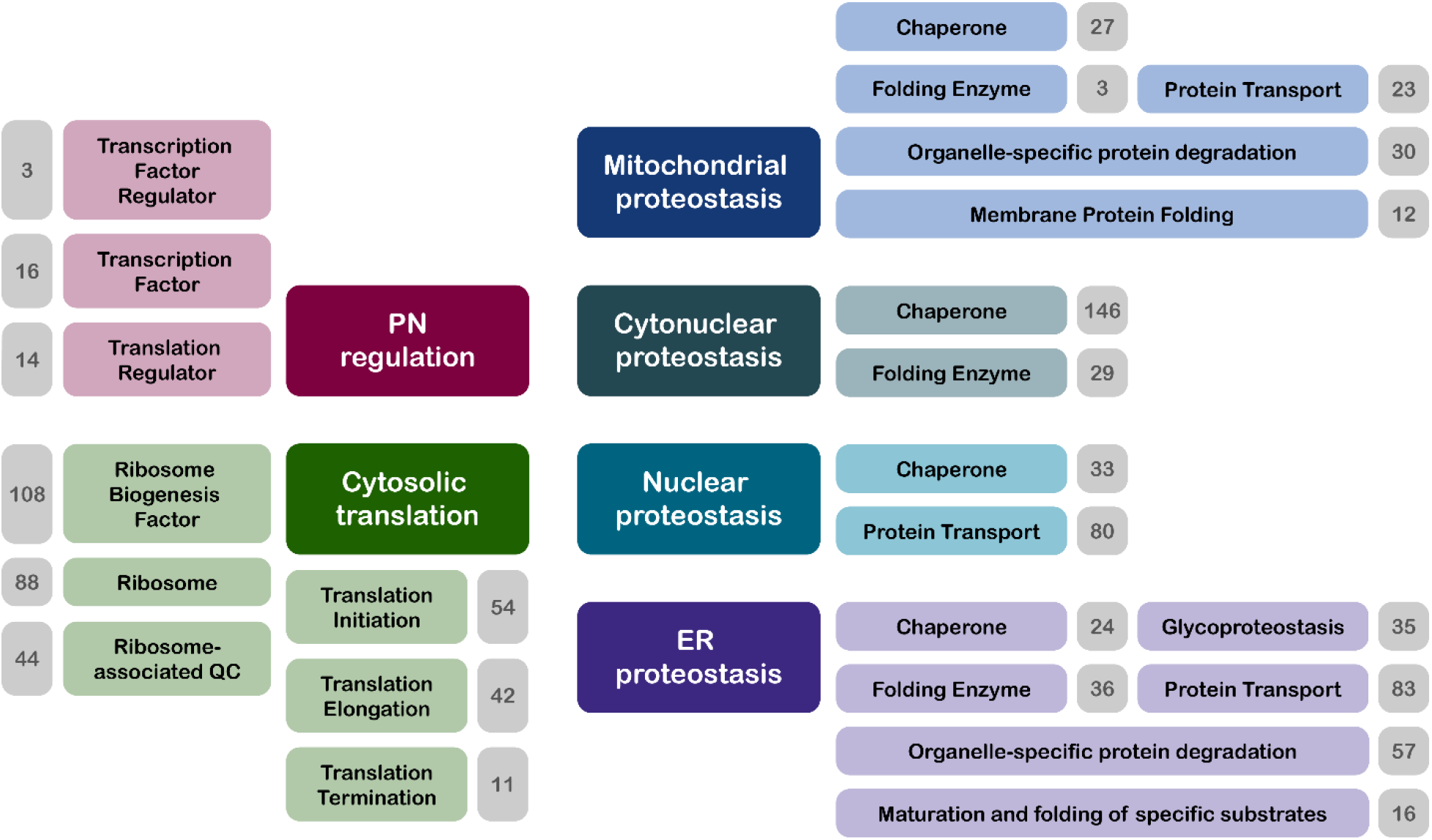
The six Branches of the proteostasis network covered in this manuscript with their cognate Classes. The numbers in the gray boxes correspond to the number of components in each class. The Classes each partition further into Groups, and in some cases, those are elaborated as Types and Subtypes. The complete hierarchy is available in Supplemental Table 1.

All 1,014 entries also have Group annotations, divided among 79 categories. A minority of entries (446, encompassing 432 components) have Type annotations, divided among 47 categories, while 153 entries (150 components) have Subtype annotations divided among 25 categories. To demonstrate how Group, Type, and Subtype annotations were assigned, we show how the cytonuclear chaperones are distributed within these finer-grained categories in Table 1. The cytonuclear chaperones (Branch = cytonuclear proteostasis, Class = chaperones) are divided into four Groups that are structurally and mechanistically distinct: the HSP70 system, the CCT/TRiC system, the sHSP (small heat shock protein) system, and the HSP90 system. Each of these Groups is then divided into the Types of components that exist within the Group. Thus, for example, there are three Types in the HSP70 system Group (Mayer and Gierasch, 2019): the HSP70 chaperones themselves, the J-domain containing cochaperones, and the HSP70 nucleotide exchange factors. Similarly, the CCT/TRiC, sHSP, and HSP90 system Groups contain three, two, and two Types respectively (Table 1). In all three of these Groups, one of the Types contains the Group’s parent chaperone (CCT/TRiC subunits, sHSPs, or HSP90). The other Types contain cochaperones, other accessory proteins, or in the CCT/TRiC system Group, components that have sequence similarity to the parent chaperone but are not part of the CCT/TRiC complex. This last Type (“sequence similar non-CCT/TRiC subunit”) was included in the proteostasis network via the domain-based criterion, but also because there is evidence that the three members of this Type form a complex with CCT/TRiC components that mediates assembly of the BBSome complex (Seo et al., 2010).

**Table 1.** Groups, Types, and Subtypes within the cytonuclear chaperone Class.

| Group | Type | Subtype | Members |
| --- | --- | --- | --- |
| **HSP70<sup>a</sup> system** | HSP70 | – | 9 |
|  | J-domain containing HSP70 cochaperone | – | 28 |
|  | HSP70 nucleotide exchange factor | HSP110/GRP170<sup>e</sup> subtype | 3 |
|  |  | BAG<sup>f</sup> domain sbtype | 6 |
|  |  | HSPBP1<sup>g</sup> subtype | 1 |
| **CCT/TRiC<sup>b</sup> system** | CCT/TRiC subunit | – | 10 |
|  | sequence similar non-CCT/TRiC subunit | – | 3 |
|  | CCT/TRiC cochaperone | Phosducin-like proteins | 5 |
|  |  | Tubulin folding | 5 |
|  |  | Prefoldin subunit | 9 |
| **sHSP<sup>c</sup> system** | small HSP | – | 12 |
|  | small HSP binding protein | – | 1 |
| **HSP90<sup>d</sup> system** | HSP90 | – | 2 |
|  | HSP90 cochaperone | CS<sup>h</sup> domain containing | 16 |
|  |  | TPR<sup>i</sup> domain containing | 20 |
|  |  | TPR and PPIase<sup>j</sup> domain containing | 5 |
|  |  | TPR and CS domain containing | 2 |
|  |  | no characteristic domain | 9 |

<sup>a</sup>HSP70 = heat shock protein 70. <sup>b</sup>CCT = chaperonin containing TCP1, TRiC = T-complex protein ring complex. <sup>c</sup>sHSP = small heat shock protein. <sup>d</sup>HSP90 = heat shock protein 90. <sup>e</sup>HSP110 = heat shock protein 110, GRP170 = glucose regulated protein 170. <sup>f</sup>BAG = BCL2-associated athanogene. <sup>g</sup>HSPBP1 = HSPA binding protein 1. <sup>h</sup>CS domain = CHORD and SGT1 domain; <sup>i</sup>TPR domain = tetratricopeptide repeat domain; <sup>j</sup>PPIase domain = peptidylprolyl isomerase domain

Some of the Types in Table 1 (three out of ten) can be further divided into Subtypes based on structural or functional criteria or both. The nucleotide exchange factor Type of the HSP70 system Group consists of three clear Subtypes based on their structures (Bracher and Verghese, 2015): the HSP110/GRP170 subtype, whose members consist of HSP70-like nucleotide- and substrate-binding domains fused to an α-helix bundle domain via an insertion of variable length (3 members); the BAG domain subtype, whose members contain a BAG domain (6 members); and the HSPBP1 subtype, whose members contain Armadillo domain repeats (1 member). There are three Subtypes of CCT/TRiC cochaperones, two of which are characterized by their functions— actin folding (5 members) and tubulin folding (5 members)—and one of which consists of subunits of the prefoldin complex (9 members). Finally, there are five Subtypes of HSP90 cochaperones whose members are characterized by the presence or absence of CS, TPR, and PPIase domains.

## Discussion

We have defined the boundaries of the human proteostasis network and presented a curated and annotated list of a selection of its components, including the cellular components responsible for protein synthesis, folding and trafficking, and protein degradation in organelles (ERAD and mitochondrial proteases). This corresponds to 959 genes, or just under 5% of human protein-coding genes. This level of investment by evolution into proteostasis illustrates the intricacies of the problem of maintaining proteostasis, which becomes more acute as organisms become more complex (Draceni and Pechmann, 2019; Powers and Balch, 2013). This observation will become even clearer in the next installments in this series, in which we will enumerate the components of the ALP and the UPS. Each of these degradative pathways are approximately the same size as the subset of proteostasis network genes described herein, illustrating just how finely tuned are the processes for protein turnover.

Our intent in providing this information on the human proteostasis network is for the community to be able to incorporate this genetic network—central for all biology—into their studies. With a comprehensive list, it will now be possible to assess which components of the proteostasis network are important for robustness and stress resilience in youth that lose capacity in aging. Likewise, assessments of the proteostasis network across the range of human diseases will uncover novel insights on protective mechanisms, basis of failure, and adaptive mechanisms. With a more complete list of the proteostasis network, it is now possible to delve deeper into Alzheimer’s diseases and related dementias and other neurodegenerative diseases to determine how and when protein quality control fails. This, in turn, will better identify potential targets for detection and therapeutic approaches.

Furthermore, we hope to facilitate the identification of systems and processes that were not known previously to engage the proteostasis network by enabling others to view their analytical frameworks through the lens of proteostasis. For example, the overlaps between a list of hits from a biochemical or bioinformatic screen and the proteostasis network could inform researchers from other fields on the contribution (or lack thereof) of proteostasis network members to the process that is the subject of the screen. Finally, we also hope to obtain feedback regarding the composition of the proteostasis network which is also shared on the Proteostasis Consortium website (proteostasisconsortium.com). Please contact us at with suggestions for components that we have excluded that should have been included, or vice versa. We regard this list as the first version of the compilation that will evolve as new experimental data on human proteostasis becomes available.

## Methods

### Criteria for inclusion in the proteostasis network

We used two criteria to decide whether a given macromolecule should be in the proteostasis network. The first was the “entity-based” criterion: an “entity” (an individual protein or non-coding RNA, or a complex) was included as a member of the proteostasis network simply if there was specific evidence in the literature that it has a role in proteostasis. The second was the “domain-based” criterion: a protein was included as a member of the proteostasis network even if there was no literature evidence that it had a role in proteostasis if it contained a domain or domains that were otherwise very strongly associated with proteostasis. This latter criterion was important for the inclusion of uncharacterized proteins that are clearly members of structural families that usually have roles in proteostasis. For example, HSPA12A and HSPA12B have not, to our knowledge, been shown to have chaperone activity, yet by sequence homology they are clearly members of the HSP70 family of chaperones (Brocchieri et al., 2008) and were therefore included as members of the proteostasis network. We show in Table 2 the literature sources that we primarily used to build our preliminary lists of proteostasis network components for the various systems mentioned above. We also used annotation databases like the Gene Ontology Resource (Consortium, 2021), the Reactome Pathway Database (Jassal et al., 2020), and the Kyoto Encyclopedia of Genes and Genomes (KEGG) (Kanehisa et al., 2016), as well as more general databases like UniProt (UniProt, 2021) and InterPro (Hunter et al., 2009) to supplement the information from the references in Table 2. The preliminary lists generated from these sources were then vetted gene by gene by the authors, with members of the Proteostasis Consortium with subject-area expertise presenting rationales for inclusion or exclusion of genes that were borderline cases. Final decisions on whether to include or exclude genes were made based on the consensus of the group.

**Table 2.**
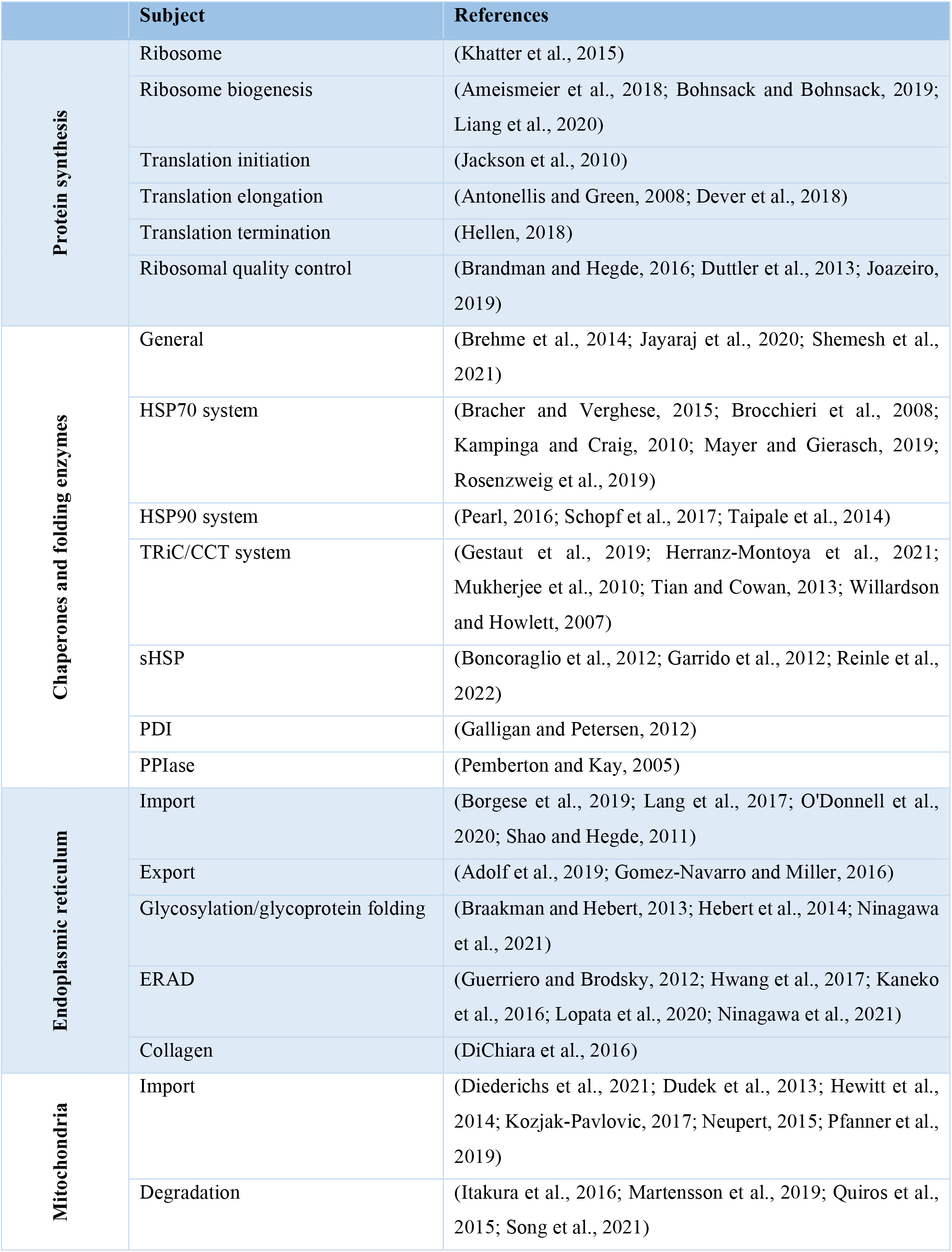
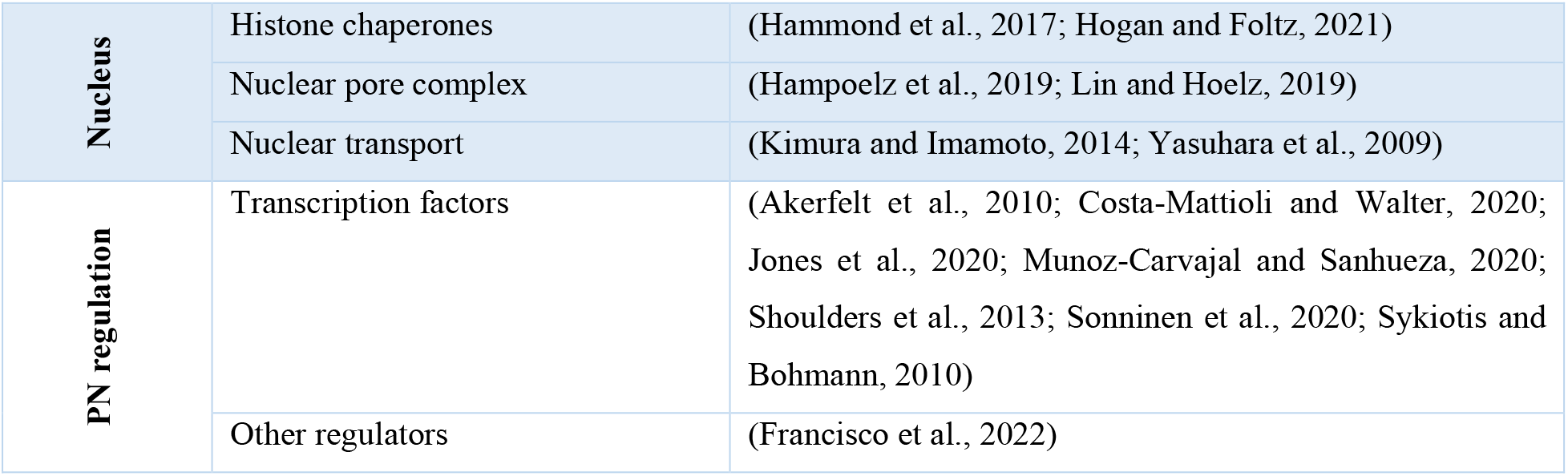
Primary literature sources for proteostasis network components.

## Acknowledgements

We thank Prof. R. Luke Wiseman (The Scripps Research Institute) for advice and feedback on the composition of the ER and mitochondrial proteostasis Branches. We gratefully acknowledge funding from the National Institutes of Health (National Institute on Aging P01AG054407 to R. I. M., D. F., S. F., J. E. G., E. T. P., J. W. K., and J. F. and K99AG068544 to T. S.).

